# Non-cognate Phosphorylation of PhoP by YycG Links Carbon Catabolite Repression to Phosphate Starvation Response

**DOI:** 10.1101/2025.06.29.661413

**Authors:** Jae-Yong Park, Wael R Abdel-Fattah

## Abstract

The PhoP-PhoR two-component system regulates transcription of the Pho regulon to counteract inorganic phosphate (Pi) limitation. Interestingly, in a *ccpA* mutant lacking the global regulator of carbon catabolism, Pho regulon genes are hyper-induced in a PhoR-independent, glucose-dependent manner.

While CcpA is known to directly repress the *phoP* promoter, we now show that overexpression of a phosphorylatable form of PhoP is essential for this hyper-induction. Remarkably, we detected early PhoP phosphorylation in the *ccpA phoR* double mutant. PhosTag-based analysis revealed that the phosphorylated/unphosphorylated PhoP ratio, rather than absolute levels, is the critical determinant of Pho regulon activation under these conditions. Given that PhoR can phosphorylate non-cognate YycF, the response regulator of the essential YycFG system controlling cell wall metabolism. We hypothesized that PhoP phosphorylation in the absence of PhoR may be mediated by YycG. Supporting this, PhoP co-immunoprecipitated with YycG, and *in vitro* phosphorylation assays confirmed direct phosphorylation of PhoP by YycG. We propose a Pi-stress regulatory model in which CcpA prevents excessive PhoP accumulation, thereby maintaining an appropriate balance between phosphorylated and unphosphorylated PhoP. This regulatory balance prevents aberrant PhoP activation via non-cognate kinases such as YycG in the absence of PhoR. Our findings highlight the interplay between carbon catabolite repression and phosphate starvation responses, and reveal a novel mechanism by which non-cognate phosphorylation contributes to adaptive transcriptional regulation.

## Introduction

*Bacillus subtilis*, a typical Gram-positive bacterium that plays an important role in soybean fermented foods such as doenjang and cheonggukjang, is often placed in natural environments that lack inorganic phosphate (Pi), such as soil. Phosphate becomes a critical limiting factor for the growth of *B. subtilis*. When faced with phosphate deficiency stress, it triggers a stress response system that controls the expression of several genes to overcome it (Hulett, 2001). The Pho regulon is regulated by the PhoP-PhoR two-component system (TCS) (Allenby et al., 2005). During periods of phosphate starvation, *B. subtilis* recognizes phosphate starvation signals through the amino-terminal sensor domain of histidine kinase (HK) PhoR, and the perceived signal is transmitted into the cell. At this time, the 360th histidine residue of PhoR is autophosphorylated. Phosphorylated PhoR (PhoR∼P) transfers phosphate to the 53rd asparagine residue of response regulator (RR) PhoP, which increases transcription of the *phoPR* gene that encodes it, as well as regulates transcription of a series of genes necessary to overcome phosphate deficiency (Hulett, 2001). These genes are about 20 or more genes or operons, including *phoA* (Ogura et al., 2001) and *phoB* (Antelmann et al., 2000), which encode alkaline phosphatase (APase), and *phoD* (Eder et al., 1996), which encodes phosphodiesterase.

*B. subtilis* is known to regulate the transcription of the *phoPR* operon by a complex regulatory apparatus, and six promoters have been found upstream of the *phoP* gene. The timing and amount of transcription at each of these promoters varies depending on the growth stage and environment (Paul et al., 2004; Puri-Taneja et al., 2006). Transcription of the *phoPR* operon requires the holoenzymes of different forms of RNA polymerase corresponding to each promoter: three promoters corresponding to σ^A^ (P_A3_, P_A4,_ and P_A6_), one corresponding to σ^B^ (P_B1_), and one corresponding to σ^E^ (P_E2_) (Kaushal et al., 2010; Paul et al., 2004; Puri-Taneja et al., 2006). For the fifth promoter, P_5_, transcription occurred only in the *sigB* mutant strain, but it is not yet known which sigma factor is required (Paul et al., 2004). The sixth and final promoter, P_A6_, was identified in the absence of CcpA, which regulates a global carbon catabolite response, and only in the *cre* (catabolite responsive element) mutant, a sequence to which CcpA binds, suggesting that carbon catabolite repression (CCR) and Pho regulon are interconnected (Puri-Taneja et al., 2006). Furthermore, transcription of four of the six promoters (P_B1_, P_E2_, P_A4,_ and P_A6_) was found to be repressed by ScoC, one of the regulators of the vegetative cell to the endospore transition state (Kaushal et al., 2010).

The phosphate starvation responses regulated by PhoPR TCS in *B. subtilis* are linked by an interdependent regulatory network involving multiple TCSs. This allows *B. subtilis* to fine-tune its responses for optimal performance under complex environmental conditions. The PhoPR TCS is part of a complex signal transduction network that includes at least three TCSs (PhoPR, ResDE, Spo0A phosphorelay), the transition state regulators AbrB and ScoC, and the CCR regulator CcpA (G. F. Sun et al., 1996). The Spo0A phosphorelay is essential for the early stationary phase and sporulation, while ResDE is involved in respiratory regulation in the late growth phase (Nakano & Hulett, 1997; Nakano et al., 2000; G. Sun et al., 1996; Zhang & Hulett, 2000).

Because the HKs and RRs that make up a TCS share very similar domains with HKs and RRs in other TCSs, there is a possibility of crosstalk with other TCSs. Although such crosstalk has been widely reported in vitro, there are few examples of this phenomenon in cells, as crosstalk between TCSs of different pathways is usually prevented by the dephosphorylating activity of HK and the competition for phosphate (Laub & Goulian, 2007). However, when bacteria allow this crosstalk under certain conditions, it can be a beneficial phenomenon, such as integrating multiple signals or differentiating one signal, called cross-regulation (Wanner, 1992). In B subtilis, reciprocal regulation has been reported to occur between PhoP-PhoR and YycF-YycG (Howell et al., 2006). In the case of *yocH*, which encodes a putative autolysin protein, its expression was YycF-dependent and not PhoP-dependent, despite being increased during phosphate starvation. Furthermore, transcription of *yocH* was absent in *phoR* mutants during phosphate starvation, and phosphorylation of YycF by PhoR was observed in vitro, suggesting that *yocH* expression is regulated by cross-regulation of YycF by PhoR during phosphate starvation (Howell et al., 2006).

This study shows the specialized conditions under which phosphorylation of PhoP by YycG can occur, in contrast to the results of Howell et al. (2006), who reported phosphorylation of YycF by PhoR. These results may explain why there is no crosstalk between YycG-induced phosphorylation of PhoP during normal phosphate-rich conditions.

## Material and methods

### Strains and plasmids

Table 1 lists the strains, plasmids, and primers used in this study. Bacterial stocks maintained at - 80°C were used to inoculate Luria-Bertani (LB) agar plates for *Escherichia coli* strains or tryptose blood agar base containing 0.5% glucose for *B. subtilis* strains. The media were supplemented with selective antibiotics at the following concentrations: 1 mg/liter; chloramphenicol (Cm), tetracycline (Tet), 10 mg/liter; kanamycin (Km), 10 mg/ml; spectinomycin (Spc), 100mg/liter; erythromycin (Em), 5 mg/liter; and ampicillin (Amp), 200 mg/liter.

**Table 1.**
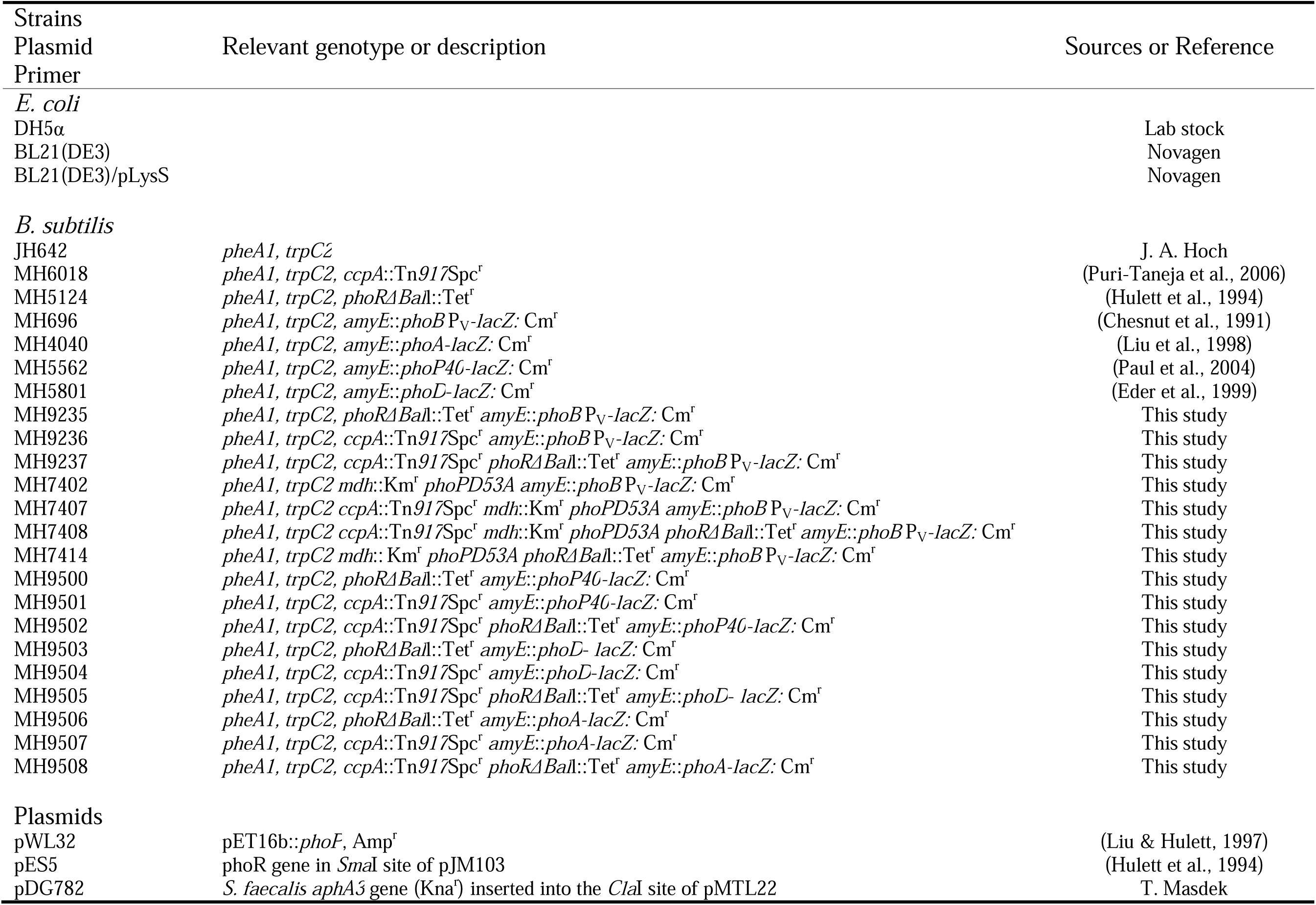
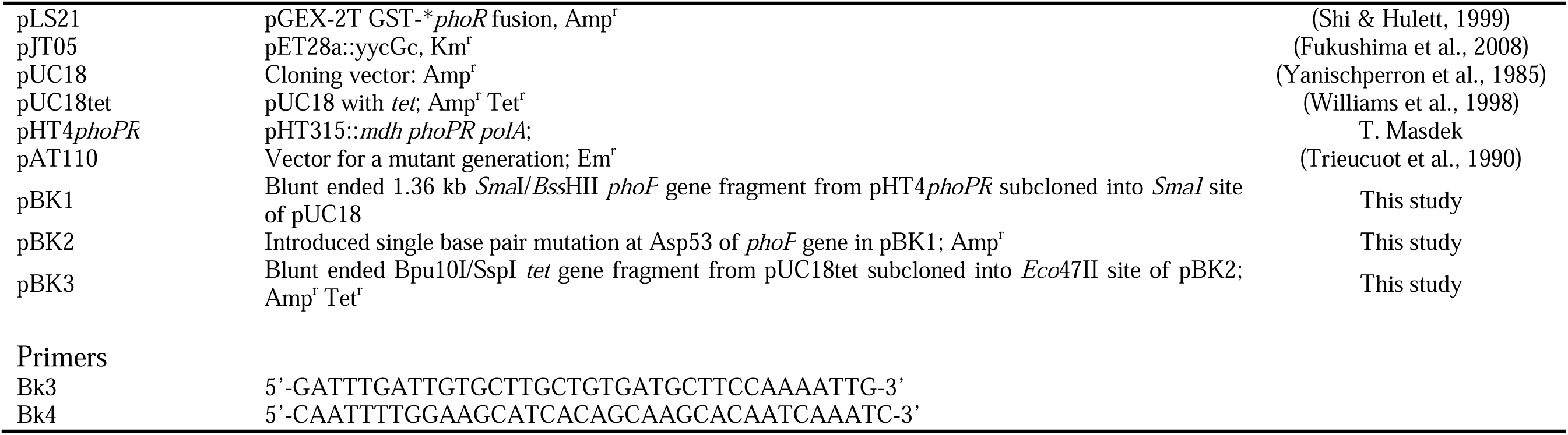

To obtain isogenic *phoR* (MH9500) or *ccpA* (MH9501) mutant strains containing *phoP40*-*lacZ* fusions, chromosomal DNA from MH5124 (Hulett et al., 1994) or MH6018 (Puri-Taneja et al., 2006) was transformed into MH5562 (Paul et al., 2004), selecting for Tet^r^ and Spc^r^ transformants respectively. An isogenic *phoR ccpA* double mutant (MH9502) with a *phoP40*-*lacZ* fusion was constructed by transforming chromosomal DNA from MH5124 (Hulett et al., 1994) into the MH9501 and selecting for Tet^r^ transformants. *phoR* (MH9235) or *ccpA* (MH9236) mutant strains, which contain a *phoB* P_V_-*lacZ* fusion, were constructed by transforming chromosomal DNA from MH5124 (Hulett et al., 1994) or MH6018 (Puri-Taneja et al., 2006) into MH696 (Chesnut et al., 1991), selecting for Tet^r^ and Spc^r^ transformants respectively. A *phoR ccpA* strain (MH9237) that contained a *phoB* P_V_-*lacZ* fusion was obtained by transforming the chromosomal DNA of MH5124 (Hulett et al., 1994) into MH9236, selecting for Tet^r^ transformants. To get *phoA*-*lacZ* fusions in *phoR* (MH9506) or *ccpA* (MH9507) mutant strains, chromosomal DNA of MH5124 (Hulett et al., 1994) or MH6018 (Puri-Taneja et al., 2006) was transformed to the MH4040 (Liu et al., 1998), selecting for Tet^r^ and Spc^r^ transformants respectively. A *phoR ccpA* strain (MH9508) with a *phoA*-*lacZ* fusion was constructed by transforming the chromosomal DNA of MH5124 (Hulett et al., 1994) to the MH9507, selecting for Tet^r^ resistant transformants. To construct isogenic *phoR* (MH9503) or *ccpA* (MH9504) that contain a *phoD*-*lacZ* fusion, chromosomal DNA of MH5124 (Hulett et al., 1994) or MH6018 (Puri-Taneja et al., 2006) were transformed to the MH5801 (Eder et al., 1999), selecting for Tet^r^ and Spc^r^ transformants respectively. A *phoR ccpA* strain containing a *phoD-lacZ* fusion (MH9505) was constructed by transforming chromosomal DNA from MH 5124 (Hulett et al., 1994) into the MH9504 and selecting for Tet^r^ transformants.

To construct non-phosphorylatable PhoPD53A strains, a 1.364 kb *Sma*I-*Bss*HII fragment from pHT4phoPR was treated with Klenow and introduced into the *Sma*I site of pUC18 (Yanischperron et al., 1985) resulting plasmid pBK1 was constructed. A single base pair mutation was inserted at Asp53 (D53A) of *phoP* by using bk3 and bk4 primer set for site-directed mutagenesis (Quick-ChangeII site-directed mutagenesis kit, Agilent, USA), resulting plasmid was named pBK2. Blunt-ended *Bpu*10I/*Ssp*I *tet* gene fragment from pUC18tet (Williams et al., 1998) was subcloned into the *Eco*47II site of pBK2; the resulting plasmid was named pBK3. Linearized pBK3 was transformed into MH696 and MH9236 to construct non-phosphorylatable PhoP_D53A_ strains, MH7402 and MH7407, respectively. To construct a *phoR* mutant, a *Bal*I deletion was made in the *phoR* gene in pES5 (Hulett et al., 1994). pDG782 (Guérout-Fleury et al., 1995), a plasmid containing the *Streptococcus faecalis aphaA3* gene as a kanamycin resistance cassette, was digested with *Sma*I/*Stu*I to release the cassette, which was cloned into the *Bal*I deletion site in pES5 resulting in pBK4. To construct a *phoR* mutation in a PhoP_D53A_ strain (MH7414), linearized pBK4 was transformed into MH7402, selecting Kan^r^ transformants. A PhoP_D53A_strain containing *ccpA* and *phoR*mutations (MH7408) was constructed by transforming linearized pBK4 into MH7407, selecting Kan^r^ transformants.

### Growth Media

Low phosphate complex medium (LPCM) plates were used to determine qualitative Pho regulon gene expression. LPCM contained 3 g/liter ammonium acetate, 0.25 g/liter MgSO_4_, 0.02 g/liter calcium acetate, 0.5 mM MnCl_2_, 10 g/liter Bacto peptone, 0.2 g/liter L-arginine, 50mM Tris (pH 6.9), 0.05 mg/ml amino acids (methionine, histidine, tryptophan, leucine, and phenylalanine), 0.05 mg/ml thiamine, and 1.5% Nobel agar (BD Difco, USA). High phosphate complex medium (HPCM, LPCM containing 10 mM K_2_HPO_4_/KH_2_PO_4_, pH 7.0), LPCMG (LPCM containing 2% glucose), HPCMG (HPCM containing 2% glucose) were also used. To assess the expression of promoter-*lacZ* fusions on solid media, X-gal (5- bromo-4-chloro-3β-D-galactosidase) was added to a final concentration of 100 mg/l.

To determine quantitative changes of total alkaline phosphatase-specific activity or *phoB* Pv-*lacZ* expression, the LPDM described previously (Hulett et al., 1990) was used with the following modifications: CoCl_2_ was omitted; ZnCl_3_ was corrected to 0.3 mM; Fructose was replaced with 2% Glucose. In growth experiments, including strains containing a *ccpA* mutation LPDMglu [low phosphate defined medium (LPDM: 0.4 mM Pi) supplemented 0.8% glutamate and 0.05 mg/ml branched-chain amino acids (BCAA)] or HPDMglu [high phosphate defined medium (HPDM: 10 mM Pi) supplemented 0.8% glutamate and 0.05 mg/ml BCAA] was used.

### Enzyme assays

For determination of total alkaline phosphatase-specific activity, 250 ml of LPDMgul or HPDMglu culture was added directly to 1 ml of the substrate, 0.1 M *p*-nitrophenyl phosphate in 1 M CHES (*N*- cyclohexyl-2-aminoethanesulfonic acid; pH 9.5), and the reaction rates were measured at an optical density at 420 nm (OD420). One unit of APase activity was equivalent to 1 μmol of *p*-nitrophenol released per min at 37°C. The APase-specific activity was expressed in units per unit of OD540.

The β-galactosidase (β-Gal) specific activity was measured by the method of Ferrari et al. (Ferrari et al., 1988) using *o*-nitrophenyl-β-D-galactopyranoside as the substrate. One unit of β-Gal activity was equivalent to 0.33 nmol of *o*-nitrophenol released per minute at 37°C. The β-Gal specific activity was expressed in units per milligram of total cellular protein. The amount of *B. subtilis* JH642 total cellular protein was calculated as previously described by Eder et al. (Eder et al., 1996).

### Overexpression and purification of proteins

For overexpression of His_10_-PhoP and GST-*PhoR (N-terminal 230 amino acids deleted PhoR) proteins, *E. coli* BL21(DE3)/pLysS was used as a host for plasmids pWL32 (Liu & Hulett, 1997), and pLS21 (Shi & Hulett, 1999), respectively (Table 1). Overexpression and purification of His-tagged and glutathione *S*-transferase (GST)-tagged proteins were performed as described previously (Liu & Hulett, 1997). *PhoR is the soluble cytoplasmic portion of the PhoR protein. For overexpression of His_6_ tagged YycG_C_ (cytoplasmic domain of YycG), *E. coli* BL21(DE3) was used as a host for plasmids pJT05 and purified as described by Fukushima et al. (Fukushima et al., 2008).

### Western blot

Western blot analysis of PhoP was performed as previously described (Abdel-Fattah et al., 2005) except for the lysis buffer [50 mM Tris-HCl (pH 7.0), 10 mM EDTA, 15 mg/ml lysozyme, 10 μg/ml DNase I, 100 μg/ml RNase A, Complete Mini EDTA-free protease inhibitor cocktail (Roche)]. After transfer, gels were fixed with Towin buffer (20% (v/v) methanol, 50 mM Tris, 40 mM glycine) and transferred to PVDF membrane at 4 a constant 100 V by using a Trans-Blot cell transfer apparatus (Bio-Rad). Antibodies were used at 1:4,000 dilutions for rabbit anti-PhoP_CTD_ antibodies (Chen et al., 2003), and secondary horseradish peroxidase-labeled antibodies were used at 1:25,000 dilutions. Bands were visualized using SuperSignal West Pico Chemiluminescent Substrate^TM^ (Pierce Biotechnology, Inc., Rockford, IL) and exposure to X-ray film.

### In vitro PhoP phosphorylation test

For confirming Phos-tag™ (Fujifilm, Japan) acrylamide SDS-PAGE system can separate phosphorylated PhoP and unphosphorylated PhoP, in vitro phosphotransfer reaction between GST-*PhoR and His_10_PhoP was performed by using γ-^32^P ATP. For autophosphorylation of GST-*PhoR, 400 nM of purified GST-*PhoR was incubated with 100 nM γ-^32^P ATP (specific activity, 6,000 Ci/mmol; 10 mCi/ml; PerkinElmer) in P buffer (50 mM HEPES [pH 8.0], 50 mM KCl, 50 mM MgCl2) at room temperature for 30 minutes. After excess γ-^32^P ATP was removed using PD SpinTrap^TM^ G-25 (GE Healthcare, Madison, USA), 400 nM of His_10_-PhoP was added for the phosphotransfer reaction. Phos-tag^TM^ acrylamide SDS-PAGE gel was prepared, described by Barbieri and Stock (Barbieri & Stock, 2008); detailed methods are described below. Phos-tag^TM^ acrylamide running gels contained 8% (w/v) 29:1 acrylamide: N, N’-methylene-bis acrylamide, 375 mM Tris-HCl, pH 8.8, 0.1% (w/v) SDS. Gels were copolymerized with 25 μM Phos-tag acrylamide and 50 μM MnCl_2_. Stacking gels contained 4% (w/v) 29:1 acrylamide: N, N’-methylene-bis acrylamide, 125 mM Tris-HCl, pH 6.8, 0.1% (w/v) SDS. Samples were mixed with 4X loading buffer (250 mM Tris pH 6.8, 8% SDS, 4% glycerol, 0.4% bromphenol blue, and 0.04% 2-mercaptoethanol) and run at 4 at a constant 15 mA/gel on 25 μM Phos- tag™ acrylamide gels until the dye front ran off the gel. The gel was fixed for 10 min in Towin buffer [20% (v/v) methanol, 50 mM Tris pH 6.8, 40 mM glycine, pH 8.3] containing 1 mM EDTA to remove Mn^2+^ from the gel. The gel was incubated for an additional 20 min in Towin buffer without EDTA. Before transfer to the PVDF membrane, the gel was exposed to x-ray film for 1 min to detect the ^32^P signal. After protein transfer to the PVDF membrane, the membrane was exposed to x-ray film for 1 min. Unphosphorylated PhoP and phosphorylated PhoP were detected as described in the western blot section.

### In vivo PhoP phosphorylation test

To detect phosphorylated PhoP in vivo, MH696, and MH9237 cells were grown in LPDMglu medium. Cells were harvested periodically by centrifugation of a 1 ml of cells adjusted to 0.5 OD_540_. The pellet was suspended with RNAlater Solution (Ambion, USA) and stored at 4 overnight before Phos-tag acrylamide gel analysis was performed. Cells were lysed with 15 μl of lysis buffer [50 mM Tris-HCl (pH 7.0), 15 mg/ml lysozyme, 10 μg/ml DNase I, 100 μg/ml RNase A, Complete Mini EDTA-free protease inhibitor cocktail (Roche, Germany), PhosSTOP Phosphatase inhibitor cocktail (Roche, Germany)] at room temperature for 3 min and solubilized by addition of 5 μl of 20% SDS and 4X loading buffer. Samples were stored on ice for a short time (<10 min) before loading onto 25 μM Phos- tag™ acrylamide gels, which were run at 4, at a constant 15 mA/gel until the dye front ran off the gel. Gel fixing, transfer, and detection were performed as described in the previous section. X-ray films were scanned using a computer scanner (Microtek ScanMaker 5), and bands were quantified using the volume report function of the ImageJ software (NIH, USA).

### Co-immunoprecipitation

MH9237 (*phoR ccpA*) cells were grown in 1 L of HPDMglu at 37°C until A600 = 0.5. Cells were harvested at 10,000 x g for 5 min and fixed with a formaldehyde solution (1% final concentration) in 30 ml P-buffer [50 mM HEPES, 50 mM KCl, 5 mM MgCl_2_ (pH 8.0)] for 20–30 min. After two washings with 50 ml phosphate buffered saline (PBS), cells were suspended in 10 ml IP buffer [PBS containing 10 mg/ml of lysozyme, 0.1 M EDTA, and EDTA-free protease inhibitor cocktail (Roche, Germany)]. The cell suspension was incubated at 37°C for 1 h before cells were lysed using the French Press G-M® (GlenMills, USA). After centrifugation at 20,000 × g for 15 min, the resulting supernatant was saved as the cytoplasmic fraction. The pellet fraction was suspended in 10 ml membrane-fraction IP buffer [PBS containing 1% Triton X-100, 0.1 M EDTA, and EDTA-free protease inhibitor cocktail (Roche)], shaken gently at room temperature for 10 min and centrifuged at 20,000 x g for 15 min. The supernatant was saved as the solubilized membrane extract. Co-immunoprecipitation was performed using an Anti- PhoP_CTD_ antibody and Pierce Co-Immunoprecipitation Kit (ThermoFisher Scientific, USA).

### In vitro phosphotransfer reaction between PhoR and PhoP or YycG and PhoP

Autophosphorylation of HKs and purification of phosphorylated HKs were performed as described in the ‘in vitro PhoP phosphorylation test’ section. 400 nM of PhoP was added to phosphorylated YycGc and GST-*PhoR, respectively. Samples were removed at different times (see Fig. 8) and mixed with a 5×SDS loading buffer (0.2 volume) to terminate the reaction. Samples were applied to 12% SDS–PAGE gels. After electrophoresis, gels were dried, and radioactivity was determined with a PhosphorImager (Azure Biosystems, USA).

## Results and discussion

### PhoR is unnecessary for expressing the Pho regulon gene in a *ccpA* mutant background

Previous studies showed that phoB PV-lacZ fusion expression in LPCM required a preferred carbon source such as glucose (LPCM) (Qi & 1998). Further, in the same medium, the *phoB* P_V_-*lacZ* fusion was expressed in a *phoR ccpA* mutant in a PhoP-dependent manner despite the absence of cognate HK PhoR (Abdel- Fattah, 2007). To determine if the expression of Pho regulon genes was common in a *phoR ccpA* mutant, we tested other Pho regulon gene expression on low or high phosphate complex medium without (LPCM; HPCM) with glucose (LPCMG; HPCMG) using *lacZ* reporter promoter fusions (Fig 1). *phoB* P_V_- (Fig 1B), *phoA*- (Fig 1C), and *phoD*-*lacZ* fusion (Fig 1D) showed similar expression patterns in each genetic background; wild type, *phoR*, *ccpA,* or *phoR ccpA* mutant strains. On LPCMG plates, *phoA*-, *phoB* P_V_-, and *phoD*-*lacZ* fusion showed high activity in wild type, *ccpA,* and *phoR ccpA* strains compared to *phoR* mutant, which showed low or no activity, indicating that the cognate HK, PhoR, was not required for Pho regulon gene induction in the *ccpA* background. These three *lacZ* fusions showed minimal expression levels on LPCM independent of strain background consistent with the requirement for a preferred carbon source for Pho induction (Qi & 1998). The data present here agree with the results of Choi and Sailer (Choi & Saier, 2005). Surprisingly, *phoA*-, *phoB* P_V_-, and *phoD*-*lacZ* fusions showed high expression levels in the *phoR ccpA* double mutant on HPCMG, while no expression was observed in other genetic backgrounds. None of the three Pho regulon promoter *lacZ* fusion strains showed expression in any genetic background on HPCM. These results suggest that Pi concentration is independent of the PhoR-independent Pho regulon induction in the *ccpA phoR* background, provided a carbon source such as glucose is present.

**Fig. 1.**
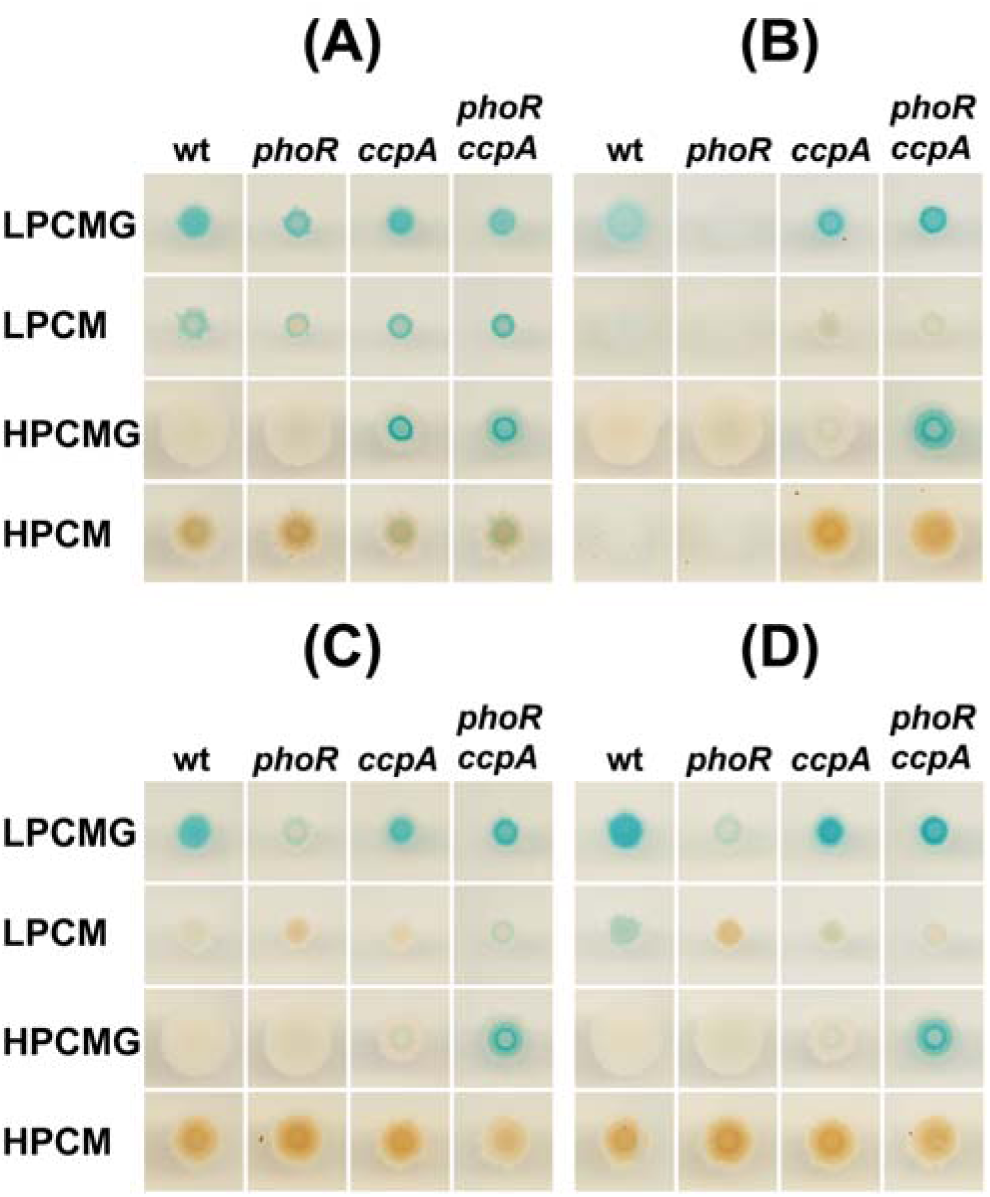
Pho regulon gene expression in wild-type and various mutant backgrounds. (A) Expression of full-length *phoP* promoter*-lacZ* fusion (*phoP40-lacZ*). (B) Expression of *phoB* P_V_*-lacZ* fusion; *phoB* (formerly *phoAIII*) encodes APaseB (formerly APaseIII). P_v_ is the σ^A^ responsive promoter of *phoB* (vegetative promoter). (C) Expression of *phoA-lacZ* fusion; *phoA* (formerly *phoAIV*) encodes APaseA (formerly APaseIV). (D) Expression of *phoD-lacZ* fusion; *phoD-lacZ* encodes a phosphodiesterase gene, PhoD. Two μl of a cell suspension containing 1 × 10^7^ cells of wild type or *B. subtilis* mutant strains carrying Pho regulon reporter fusions (*phoB* Pv, *phoA,* or *phoD*) or the auto-regulated *phoP* fusion were applied to the surface of LPCMG (low phosphate complex medium with glucose), LPCM (low phosphate complex medium without glucose), HPCMG (high phosphate complex medium with glucose), and HPCM (high phosphate complex medium without glucose) plates containing X-Gal. Plates were incubated at 37℃ for 48 hours.

The *phoP*40-*lacZ* fusion strains (Fig 1A) showed similar expression patterns in each genetic background on LPCMG or LPCM, with the *phoR* mutant showing the lowest expression level on either medium. The *phoP*40-*lacZ* fusion showed a dramatic increase of expression in the *ccpA* and *phoR ccpA* double mutant backgrounds grown on HPCMG compared to wild-type and *phoR* mutants that showed no expression. This is consistent with previous studies showing that in the presence of a carbon source such as glucose, the *ccpA* mutation releases repression of the *phoPR* P_A6_ promoter independent of Pi concentration (Puri-Taneja et al., 2006).

### Pho regulon gene induction depends not on the quantity of PhoP but on the phosphorylation of PhoP

Because either phosphorylated PhoP or non-phosphorylated PhoP can transcribe Pho regulon genes in vitro, we reasoned that PhoP concentrations in a *ccpA* mutant strain might be high enough to activate Pho regulon genes in an unphosphorylated state. In vitro, transcription studies using the *phoBPv* promoter had shown that 16-fold more non-phosphorylated PhoP than phosphorylated PhoP was required to transcribe 85% of the maximal transcription yield of phosphorylated PhoP (Abdel-Fattah et al., 2005). Further, in vivo expression of the *phoPR* promoter fusion in a *ccpA* mutant strain was increased three or morefold (depending on the growth phase) compared to a WT strain. This expression resulted from the derepression of PA6, which could not be detected in a WT strain (Puri-Taneja et al., 2006). To determine if the PhoP concentration in the *ccpA* mutant is higher than the wild-type strain, we carried out a quantitative Western blot analysis of PhoP synthesis during growth in a low phosphate defined medium with glutamate and branched-chain amino acids (LPDMglu). The growth defect of a *ccpA* mutant in the defined medium is well understood; CcpA is necessary for the transcription of *liv-leu* operon for the biosynthesis of branched-chain amino acids (BCAA) and *gltAB* encoding glutamate synthase (Sonenshein, 2007). To circumvent the *ccpA* mutant growth defect, we added BCAA (50 ug/ml) and glutamate (final 0.8%) in LPDM (Ludwig et al., 2002). The results showed that PhoP concentrations were increased in a *ccpA* mutant compared to the wild-type strain (Fig. 2), and the fold increase depended on the growth phase.

**Fig. 2.**
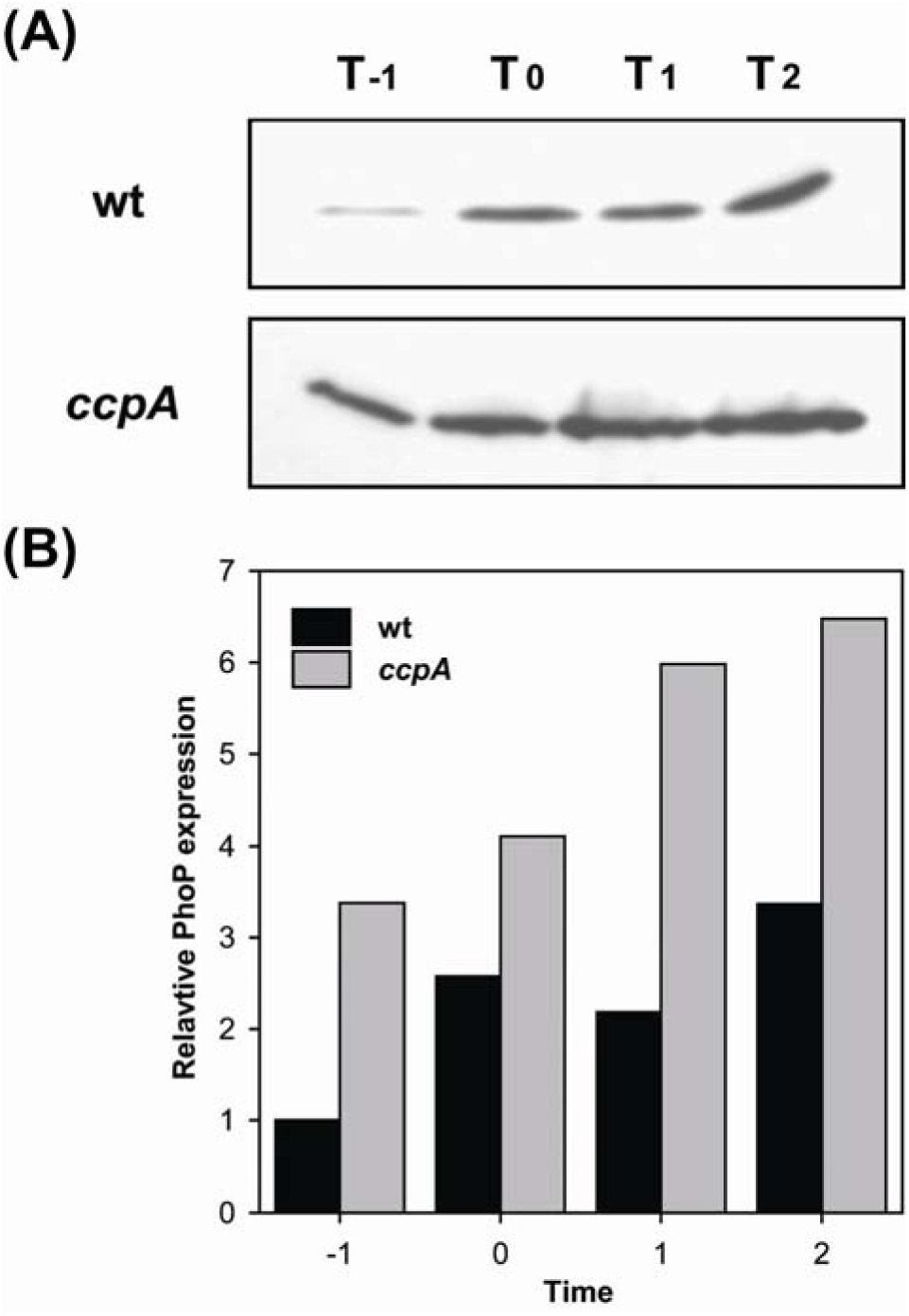
Expression of PhoP in wild-type strain and *ccpA* mutant in LPDMglu. (A) Western blot detection of PhoP in wild-type strain and *ccpA* mutant. (B) Relative PhoP expression level based on PhoP expression level of wild type at T_-1_. For Western blot detection of intracellular PhoP concentration, samples were collected at indicated times (T_0_ means transition state). After microscopic cell counting, 5 × 10^7^ cells were used for western blot as described in Materials and Methods.

To test the hypothesis that high concentrations of non-phosphorylated PhoP were responsible for the expression of Pho genes in a *ccpA phoR* double mutant, we needed to determine if Pho regulon gene expression occurs in strains having only a non-phosphorylatable PhoP. pBK3, a vector for replacing *phoP* to *phoP*_D53A_ encoding non-phosphorylatable PhoP, was constructed as mentioned in the materials and methods. We introduced PhoP_D53A_ mutation in the wild type, *phoR*, *ccpA*, and *phoR ccpA* strains. *phoB* P_V_-*lacZ* fusion expression was tested on complex media plates, LPCMG and HPCMG (Fig. 3A), or defined media LPDM and HPDM plates (Fig. 3B) containing glutamate and BCAA where required for growth. All PhoP_D53A_ mutant strains failed to induce *phoB* P_V_-*lacZ* fusion on plates containing each of these media (Fig. 3AB), while controls containing a wild-type *phoP* gene expressed as expected based on data from Fig. 1. We also tested Pho induction of PhoP_D53A_ mutant strains in LPDMglu liquid medium (containing BCAA and glutamate) by measuring total APase specific activity as Pho induction reporter. All PhoP_D53A_ mutants grew well but showed no PhoP induction (Fig. 3C), suggesting phosphorylated PhoP is necessary for Pho regulon induction. Together, these data indicated that the Pho regulon expression in the *phoR ccpA* mutant strain depended not on the PhoP amount but on PhoP phosphorylation.

**Fig. 3.**
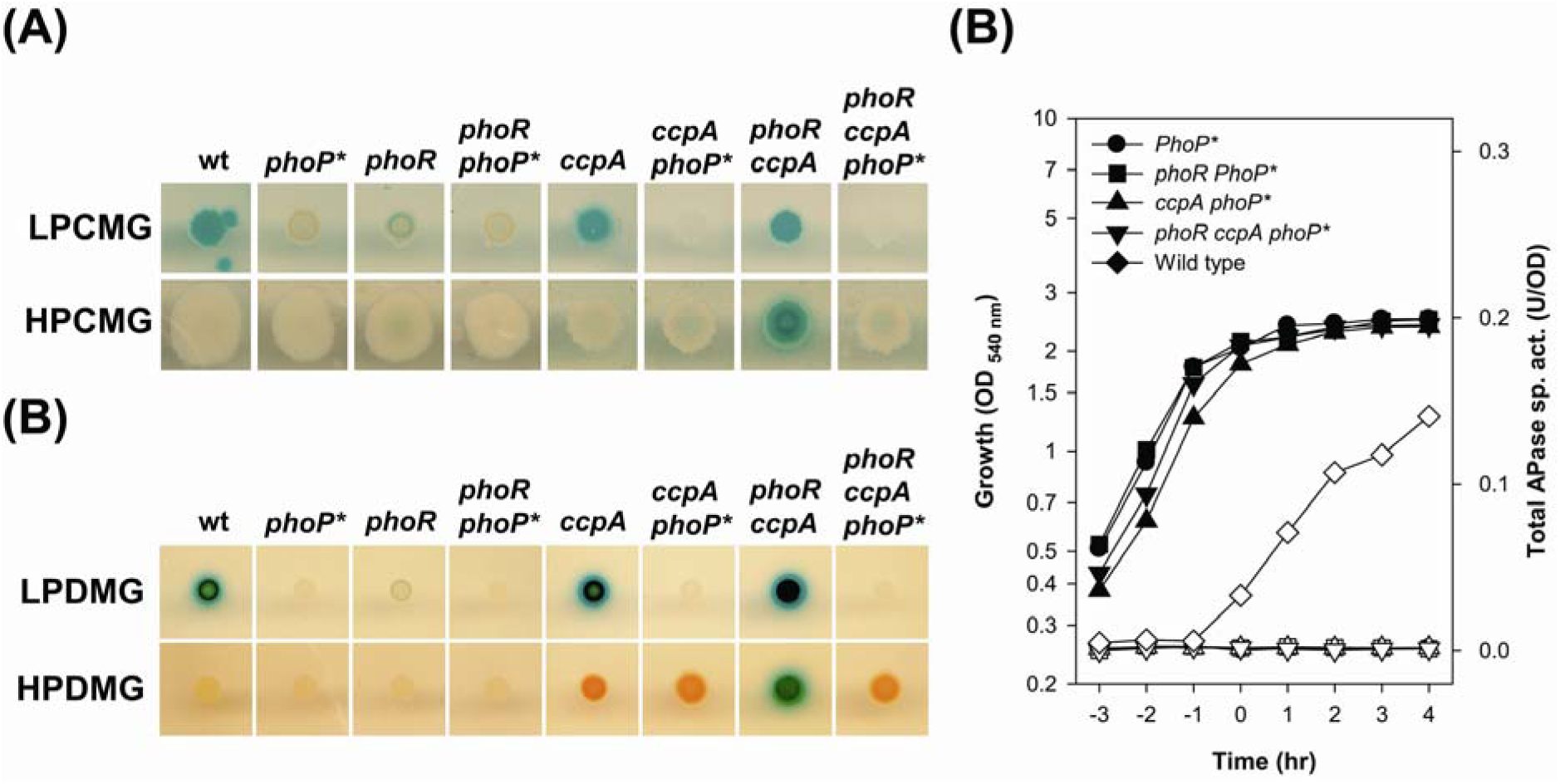
*phoB* P_V_-*lacZ* expression in various strains containing non-phosphorylatable PhoPD53A. (A) *phoB* P_V_-*lacZ* expression on LPCMG and HPCMG plates. (B) *phoB* P_V_-*lacZ* expression on LPDM and HPDM plates, low or high phosphate defined medium containing glutamate and BCAA as required for growth. *phoP*\* indicates a *phoP*D53A strain. *B. subtilis* strains were inoculated and grown as described in Fig 1. (C) Growth curve (closed symbols) and total APase specific activities (open symbols) of strains containing non-phosphorylatable PhoPD53A in LPDMGglu. Circles, squares, triangles, and inverted triangles indicate wild type, *phoR*, *ccpA,* and *phoR ccpA* mutant strains containing nonphosphorylatable PhoP(PhoP*), respectively. Diamonds indicate a wild-type strain containing the wild-type *phoP* gene as a positive control.

### The PhoR-independent Pho induction is independent of Pi concentration in the *phoR ccpA* mutant strain

To determine the temporal and quantitative expression of *phoB*P_V_-*lacZ* in the *phoR ccpA* background, we used LPDMglu and HPDMglu media. We assayed growth and *phoB* P_V_-*lacZ* expression of wild-type and *phoR ccpA* mutant strains in LPDMglu (Fig 4 A) and HPDMglu (Fig 4 B). It has been reported that when the Pi concentration of the LPDM culture decreased to 0.1 mM, the wild-type strain stopped cell division, initiating the transition phase (T_0_) and Pho induction (Abdel-Fattah et al., 2005; G. F. Sun et al., 1996). The wild-type strain showed traditional Pho induction in LPDMglu (Fig 4A -○-). The *phoR ccpA* mutant showed low-level *phoB* P_V_-*lacZ* expression constitutive throughout growth (Fig 4A -▽-). In HPDMglu, the *phoR ccpA* mutant expressed similar levels of *phoB* P_V_-*lacZ* expression observed in LPDMglu (Fig 4B -▽-), while the wild-type strain showed no Pho induction (Fig 4B -○-). Consistent with the plate assays (Fig. 1 and 3), these data indicated that PhoR-independent Pho expression in *phoR ccpA* mutant was independent of Pi concentration.

**Fig. 4.**
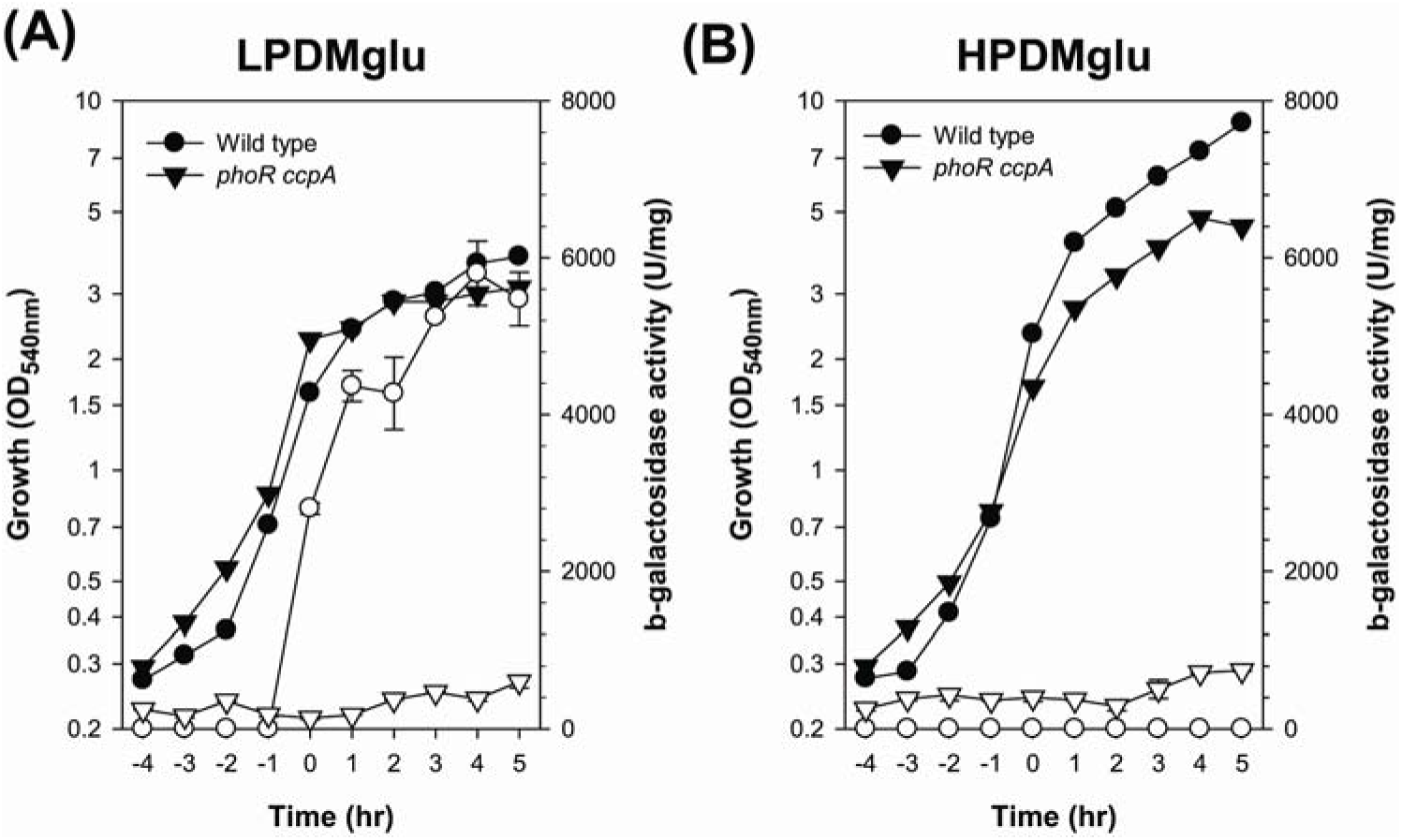
Temporal induction of *phoB* P_V_-*lacZ* in wild type and *phoR ccpA* mutant strains in LPDMglu (A) and HPDMglu (B). Closed symbols indicate growth and open symbols indicate β-Gal activities—circles (wild type): inverted triangles (*phoR ccpA* mutant).

### Pho regulon gene expression in a *phoR ccpA* mutant depended on PhoP phosphorylation

To determine the in vivo PhoP phosphorylation state in the *phoR ccpA* mutant, we employed Phos-tag™ acrylamide SDS-PAGE and Western blot techniques. Before initiating in vivo tests, we needed to determine if Phos-tag™ acrylamide SDS-PAGE can separate PhoP from PhoP∼P. We performed Phos- tag™ acrylamide SDS-PAGE after an in vitro phosphotransfer reaction between autophosphorylated GST-*PhoR∼^32^P and PhoP. The images on X-ray film exposed to the wet gel and the PVDF membrane showed autophosphorylated GST-*PhoR detected PhoP∼P signals (Fig. 5A-B). When we performed Western blot analysis using the same PVDF membrane and PhoP_CTD_-specific Ab, two signals were detected (Fig. 5C, lane 3); one which migrated with PhoP (Fig 5C lane 1) and the other PhoP∼P (Fig 5 A and 5B, lanes 3). We concluded that Western blot combined Phos-tag™ acrylamide SDS-PAGE was potentially a powerful tool for detecting PhoP∼P in vivo.

**Fig. 5.**
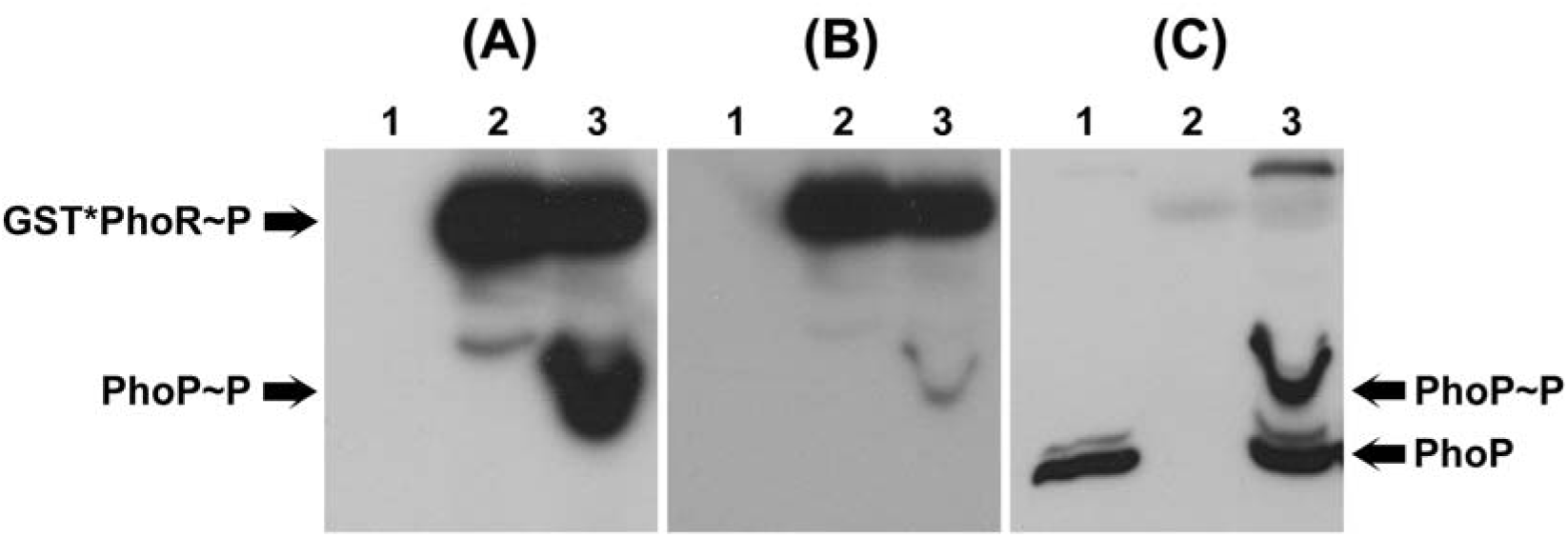
Separation of PhoP and PhoP∼P by using Phos-tag™ acrylamide SDS-PAGE and γ-^32^P ATP. After phosphotransfer from GST-*PhoR∼P to PhoP, Phos-tag™ acrylamide gel separation was performed, and the gel was exposed to an x-ray film for 18 hr (A). The proteins were transferred to the PVDF membrane, which was exposed to x-ray film for 4 hr (B). Western blot was performed on the membrane using PhoP_CTD_ specific antibody (C). Lane 1, PhoP; lane 2, autophosphorylated GST-*PhoR; lane 3, autophosphorylated *GST-PhoR plus PhoP.

To assay PhoP phosphorylation in vivo in the *phoR ccpA* mutant, wild-type and *phoR ccpA* strains were grown in LPDMGglu. APase activity was measured as a Pho induction reporter, and cells were collected for Phos-tag™ acrylamide SDS-PAGE analysis. As expected, the wild-type strain expressed APase activity only after T_0_. In contrast, the *phoR ccpA* mutant expressed an initial APase-specific activity that slowly decreased with time (Fig. 6B). In vivo PhoP∼P was detected (Fig 6A) in the wild-type strain between T_1_ and T_5_. In the *phoR ccpA* mutant strain, PhoP∼P was detected in all samples from T_-3_ to T_2_. These data confirmed that PhoP was phosphorylated in vivo in *phoR ccpA* mutant despite the absence of PhoR, presumably by another HK or small molecular phosphor donor.

**Fig. 6.**
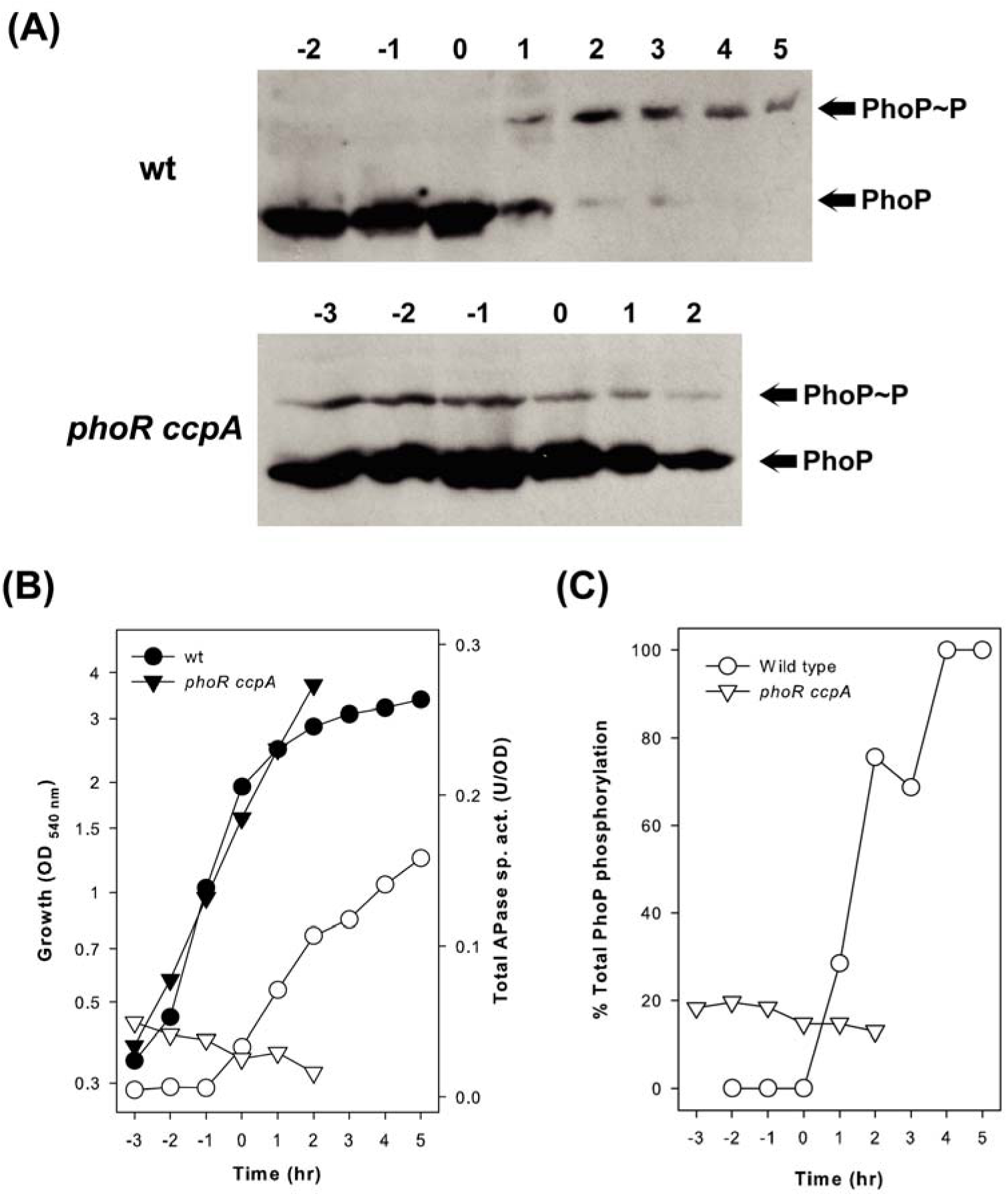
In vivo PhoP phosphorylation in wild type and *phoR ccpA* strains. (A) Detection of in vivo phosphorylated PhoP during growth in LPDMglu using Phos-tag™ acrylamide SDS-PAGE combined with Western Blot analysis. (B) Growth (close symbols) and total APase specific activities (open symbols) in LPDMglu. Circles (wild type strain); inverted triangles (*phoR ccpA* mutant). (C) PhoP∼P as % of total PhoP in wild type and *phoR ccpA* strains.

The percentage of total PhoP, which was phosphorylated (Fig 6C), showed a similar pattern to that of APase-specific activity (Fig 5B). The absolute quantity of PhoP∼P in the *phoR ccpA* mutant at T_-3_ was about 2-fold higher than the wild-type strain at T_5_. Still, the APase-specific activity was approximately 3.2-fold lower than the wild type at T_5,_ suggesting that the Pho induction level is determined by the phosphorylated proportion of total PhoP rather than the absolute amount of intracellular PhoP∼P. The fact that the % of total PhoP in the cell that was phosphorylated, rather than the level of PhoP∼P in the cell, was necessary for the level of Pho regulon gene expression was unexpected and raises the question of negative regulation by unphosphorylated PhoP in vivo. It was shown in vitro that both PhoP and PhoP∼P can bind to the PhoP binding consensus repeats in all Pho regulon genes tested, although PhoP∼P usually had a higher binding affinity. We had previously observed that *phoP* was expressed from a multicopy plasmid in a wild-type strain. In this case, Pho regulon gene induction was decreased compared to expression in a wild-type strain (Qi & 1998).

### YycG may phosphorylate PhoP in a *phoR ccpA* mutant strain

The data presented above prompted the question: what is PhoP phosphorylated by in the phoR ccpA mutant that lacks PhoR? One could hypothesize that PhoP was phosphorylated in vivo by a non-cognate HK or small molecule phosphodonor in a *phoR ccpA* strain. Intermediary metabolite phosphodonors, such as acetyl phosphate and carbamoyl phosphate, can phosphorylate *E. coli* RRs such as CheY (Lukat et al., 1992). The *E. coli* PhoP orthologue, PhoB, can also be phosphorylated by acetyl phosphate in vitro and in vivo (Kim et al., 1996; McCleary & Stock, 1994). Because PhoP phosphorylation by small molecular phosphodonors failed in vitro (Liu & Hulett, 1997), we considered another HK more likely to phosphorylate PhoP in vivo than a small-molecule phosphodonor.

Fabret et al. classified *B. subtilis* HKs based on the similarity of sequences adjacent to the phosphorylated histidine residue. They further sub-grouped the kinases based on their transmembrane segment structure (Fabret et al., 1999). ResE and YycG have similar amino acid sequences around the phosphorylated histidine residue and transmembrane structures like PhoR. Therefore, we considered these two HKs likely candidates for in vivo crosstalk partner(s) of PhoP in *phoR ccpA* mutant.

ResE was considered a crosstalk partner because expression patterns of cognate RRs showed that ResD expression increased at a similar stage of Pho induction as PhoP (Abdel-Fattah *et al*., unpublished data). Still, the exponential expression of YycF expression stopped at T_0_ (Fabret & Hoch, 1998). The possibility that ResE phosphorylated PhoP was especially attractive because the ResDE TCS is connected to the Pho signal transduction network (Birkey et al., 1998; Schau et al., 2004). ResE can phosphorylate PhoP in vivo (Abdel-Fattah *et al*., unpublished data), but wild-type, *phoR*, *ccpA*, and *phoR ccpA* strains containing a *resE* mutation did not lose *phoB* P_V_-*lacZ* activities in LPCMG plates (data not shown). Moreover, phosphorylation of PhoP in the *ccpA phoR* mutant occurs before PhoP phosphorylation or Pho induction in the WT strain (Fig. 6A&B) and before Pho or Res induction in the WT strain grown in LPDM (1; Abdel-Fattah and Hulett, unpublished data). Together, these data suggested that the phosphoryl donor for PhoP in the *phoR ccpA* strain was not dependent on ResE.

Similar experiments to determine if YycG were the PhoP phosphodoner were complicated because the YycG-YycF TCS is essential for cell growth (Fabret & Hoch, 1998). Because the highest levels of PhoP∼P were observed in *phoR ccpA* cells during non-limiting phosphate growth conditions, T_-3_ to T_-1_ (Fig. 6A), we designed experiments to determine if PhoP interacts with YycG in *phoR ccpA* mutant during non-limiting phosphate conditions. Co-immunoprecipitation with PhoP_CTD_ specific antibody was performed with MH9237 (*phoR ccpA*) cells grown in HPDMglu after formaldehyde cross- linking and cell lysis. Western blot analysis, using the co-immuno-precipitated protein sample, showed PhoP and YycG signals that superimpose at approximately 150 kDa (Fig. 7). The size of the cross- linked complex suggested that either a YycG monomer interacted with a PhoP dimer (about 125.2 kDa), trimer (about 152.8 kDa) or a YycG dimer interacted with a PhoP monomer (about 167.7 kDa). The intermediate-sized bands that cross-react with both antibodies are likely proteolytic products of the isolated complex.

**Fig. 7.**
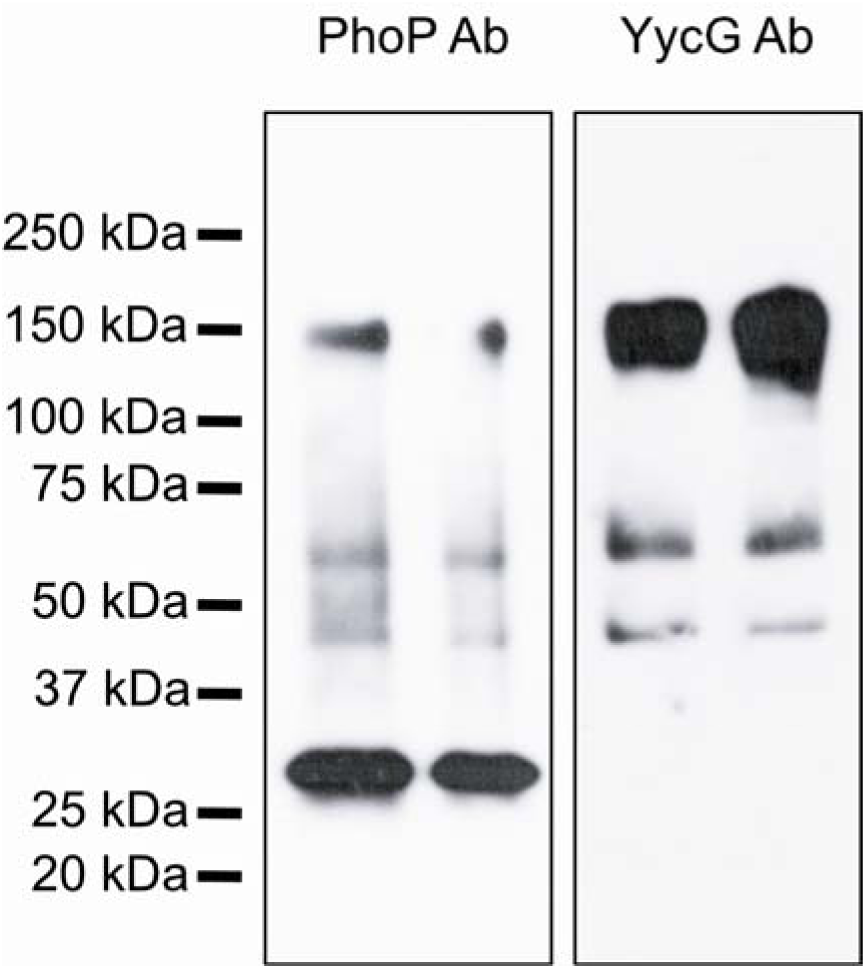
YycG co-precipitates with PhoP in *phoR ccpA* mutant in an immunoprecipitation assay. As described in Materials and Methods, exponentially growing phoR ccpA mutant cells (OD540= 0.5) were subjected to cross-linking and immunoprecipitation from cell extracts with anti-PhoPCTD antibodies. PhoP (left panel) and YycG (right panel) proteins were visualized immunologically by Western blot analysis using anti-PhoP_CTD_ or anti-YycG_C_ antibodies, respectively.

**Fig. 8.**
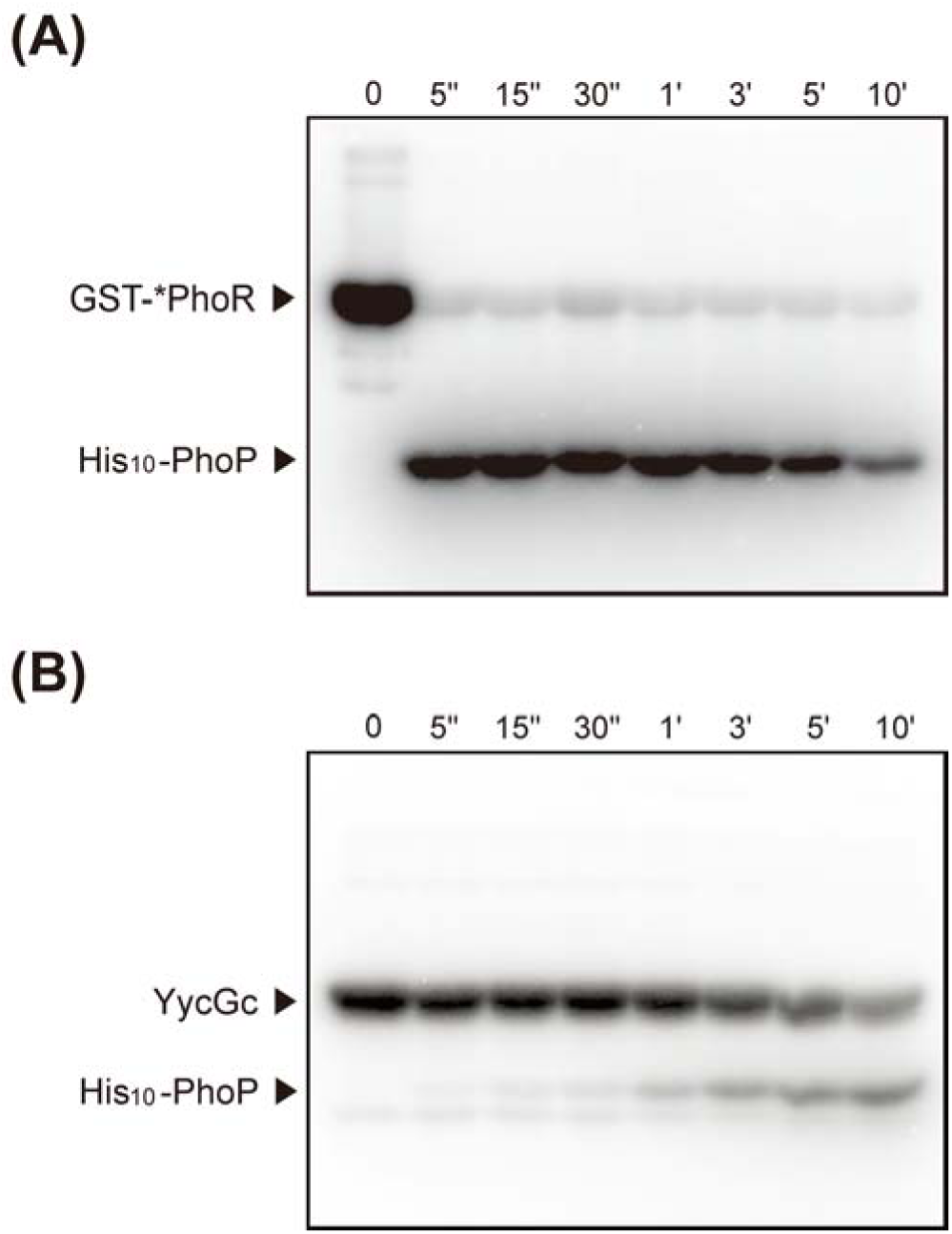
In vitro phosphotransfer between GST-*PhoR and His_10_-PhoP or YycGc and His_10_-PhoP. Samples were collected at indicated time intervals, denatured, and subjected to SDS-PAGE. Dried gels were visualized using PhosphorImager. (A) Phosphotransfer between GST-*PhoR and His_10_-PhoP. (B) Phosphotransfer between YycGc and His_10_-PhoP.

We compared the in vitro phosphotransfer between PhoR∼P and PhoP with YycG∼P and PhoP. Phosphotransfer from PhoR∼P to PhoP was complete within 5 seconds (Fig. 8A); Phosphotransfer from YycG∼P to PhoP could be detected within 5 seconds, but the rate and the efficiency of the phosphotransfer were significantly reduced (Fig. 8B) compared to that of PhoR∼P. Together, these results supported the supposition that YycG may phosphorylate the non-cognate RR PhoP in a *ccpA phoR* mutant strain but independent of limiting Pi concentrations and much less effectively than PhoR.

HK and RR composing a TCS may be a crosstalk phenomenon because HK and RR have a significantly similar domain to HK and RR comprising other TCSs. Few crosstalk phenomena are reported in vivo because cognate HK prevents it via phosphatase activity and substrate competition despite numberless in vitro reports (Laub & Goulian, 2007). Howel et al. suggested the in vitro crosstalk phenomenon between PhoP-PhoR and YycF-YycG (Howell et al., 2006). The expression of the *yocH* gene, encoding a protein-predicted autolysin, is not dependent on PhoP but depends on YycF, despite yocH induced during the phosphate starvation state. Furthermore, *yocH* does not transcribe during the phosphate starvation state in *phoR* mutant, and PhoR can phosphorylate YycF in vitro. Howel et al. conclude that crosstalk between PhoR and YycF controls *yocH* expression in phosphate deficiency conditions (Howell et al., 2006). The results presented here show that PhoP can be phosphorylated by YycG when cognate HK PhoR and carbon catabolite regulator CcpA are absent. Our in vitro phosphotransfer reaction data between YycG and PhoP argued with the data that YycG could not phosphorylated PhoP in vitro showed Howell et al. (Howell et al., 2006). We suppose the difference may be due to the different methods employed for the phosphotransfer reaction or the protein purification procedures.

In conclusion, the phosphate concentration-independent expression of Pho regulon genes in the *ccpA phoR* mutant is likely due to the phosphorylation of PhoP by YycG, a non-cognate HK. The reason for this phenomenon seems to be that the repression of PhoP expression by CcpA is released, and the accumulation of a large amount of PhoP in the cell leads to a low level of phosphorylation of PhoP by YycG, which is unlikely to occur. In other words, CcpA regulates the Pho regulon by suppressing the overexpression of PhoP, thereby preventing crosstalk that could be caused by other HKs such as YycG.

## Acknowledgments

This work was supported by research grants (20231149) from Daegu Catholic University in 2023. Jae-Yong Park was supported by the Korea Research Foundation Grant (KRF-2007-357-F00040), funded by the Korean Government (MOEHRD). We thank James A. Hoch for providing an anti-YycG antibody and YycG overexpression plasmid pJT05.

## Data availability

Data are available on request from the authors.

## Declarations Conflict of interest

The authors declare no commercial or financial conflict of interest.

## Ethical Statement

The manuscript does not contain experiments using animals or human studies.

## Notes

### Competing Interest Statement

The authors have declared no competing interest.

## References

1. Abdel-Fattah, W. R. (2007). Bacillus subtilis Pho∼P Direct Roles in Pho and Res Regulation in Response to Pi-Stress [Doctor of Philosophy., University of Illinois at Chicago ]. Chicago, Illinois.

2. Abdel-Fattah, W. R., Chen, Y. H., Eldakak, A., & Hulett, F. M. (2005). *Bacillus subtilis* phosphorylated PhoP: Direct activation of the E sigma(A)- and repression of the E sigma(E)-responsive *phoB*-PS+V promoters during Pho response [Article]. Journal of Bacteriology, 187(15), 5166–5178. 10.1128/jb.187.15.5166-5178.2005

3. Allenby, N. E., O’Connor, N., Prágai, Z., Ward, A. C., Wipat, A., & Harwood, C. R. (2005). Genome-wide transcriptional analysis of the phosphate starvation stimulon of *Bacillus subtilis*. Journal of Bacteriology, 187(23), 8063–8080. 10.1128/jb.187.23.8063-8080.2005

4. Antelmann, H., Scharf, C., & Hecker, M. (2000). Phosphate starvation-inducible proteins of *Bacillus subtilis*: proteomics and transcriptional analysis. Journal of Bacteriology, 182(16), 4478–4490. 10.1128/jb.182.16.4478-4490.2000

5. Barbieri, C. M., & Stock, A. M. (2008). Universally applicable methods for monitoring response regulator aspartate phosphoylation both in vitro and in vivo using Phos-tag-based reagents. Analytical Biochemistry, 376, 73–82.

6. Birkey, S. M., Liu, W., Zhang, X. H., Duggan, M. F., & Hulett, F. M. (1998). Pho signal transduction network reveals direct transcriptional regulation of one two-component system by another two-component regulator: *Bacillus subtilis* PhoP directly regulates production of ResD [Article]. Molecular Microbiology, 30(5), 943–953. <Go to ISI>://000077817100005

7. Chen, Y. H., Birck, C., Samama, J. P., & Hulett, F. M. (2003). Residue R113 is essential for PhoP dimerization and function: a residue buried in the asymmetric PhoP dimer interface determined in the PhoPN three-dimensional crystal structure [Article]. Journal of Bacteriology, 185(1), 262–273. 10.1128/jb.185.1.262-273.2003

8. Chesnut, R. S., Bookstein, C., & Hulett, F. M. (1991). Separate promoters direct expression of *phoAIII*, a member of the Bacillus subtilis alkaline-phosphatase multigene family, during phosphate starvation and sporulation [Article]. Molecular Microbiology, 5(9), 2181–2190. <Go to ISI>://A1991GG05100014

9. Choi, S. K., & Saier, M. H. (2005). Regulation of pho regulon gene expression by the carbon control protein A, CcpA, in *Bacillus subtilis* [Article]. Journal of Molecular Microbiology and Biotechnology, 10(1), 40–50. 10.1159/000090347

10. Eder, S., Liu, W., & Hulett, F. M. (1999). Mutational analysis of the *phoD* promoter in *Bacillus subtilis*: Implications for PhoP binding and promoter activation of Pho regulon promoters [Article]. Journal of Bacteriology, 181(7), 2017–2025. <Go to ISI>://000079368400006

11. Eder, S., Shi, L., Jensen, K., Yamane, K., & Hulett, F. M. (1996). A *Bacillus subtilis* secreted phosphodiesterase alkaline phosphatase is the product of a Pho regulon gene, phoD [Article]. Microbiology-Uk, 142, 2041–2047. <Go to ISI>://A1996VB82900016

12. Fabret, C., Feher, V. A., & Hoch, J. A. (1999). Two-component signal transduction in *Bacillus subtilis*: How one organism sees its world [Review]. Journal of Bacteriology, 181(7), 1975–1983. <Go to ISI>://000079368400001

13. Fabret, C., & Hoch, J. A. (1998). A two-component signal transduction system essential for growth of *Bacillus subtilis*: Implications for anti-infective therapy [Article]. Journal of Bacteriology, 180(23), 6375–6383. <Go to ISI>://000077217600036

14. Ferrari, E., Henner, D. J., Perego, M., & Hoch, J. A. (1988). Transcription of *Bacillus subtilis* subtilisin and expression of subtilisin in sporulation mutants [Article]. Journal of Bacteriology, 170(1), 289–295. <Go to ISI>://A1988L506300044

15. Fukushima, T., Szurmant, H., Kim, E.-J., Perego, M., & Hoch, J. A. (2008). A sensor histidine kinase co-ordinates cell wall architecture with cell division in *Bacillus subtilis*. Molecular Microbiology, 69(3), 621–632. 10.1111/j.1365-2958.2008.06308.x

16. Guérout-Fleury, A.-M., Shazand, K., Frandsen, N., & Stragier, P. (1995). Antibiotic-resistance cassettes for *Bacillus subtilis*. Gene, 167, 335–336. doi:10.1016/0378-1119(95)00652-4

17. Howell, A., Dubrac, S., Noone, D., Varughese, K. I., & Devine, K. (2006). Interactions between the YycFG and PhoPR two-component systems in *Bacillus subtilis*: the PhoR kinase phosphorylates the non-cognate YycF response regulator upon phosphate limitation [Article]. Molecular Microbiology, 59(4), 1199–1215. 10.1111/j.1365-2958.2005.05017.x

18. Hulett, F. M. (2001). The Pho Regulon. In A. L. Sonenshein, J. A. Hoch, & R. Losick (Eds.), Bacillus subtilis and Its Closest Relatives: From Genes to Cells (pp. 193–201). ASM Press. 10.1128/9781555817992.ch15

19. Hulett, F. M., Bookstein, C., & Jensen, K. (1990). Evidence for 2 structural genes for alkaline-phosphatase in *Bacillus subtilis*. Journal of Bacteriology, 172(2), 735–740. <Go to ISI>://WOS:A1990CL74300030

20. Hulett, F. M., Lee, J. W., Shi, L., Sun, G. F., Chesnut, R., Sharkova, E., . . . Kapp, N. (1994). Sequential action of two-component genetic switches regulates the PHO regulon in *Bacillus subtilis* [Article]. Journal of Bacteriology, 176(5), 1348–1358. <Go to ISI>://A1994MY35100019

21. Kaushal, B., Paul, S., & Hulett, F. M. (2010). Direct regulation of *Bacillus subtilis phoPR* transcription by transition state regulator ScoC. Journal of Bacteriology, 192(12), 3103–3113. 10.1128/jb.00089-10

22. Kim, S. K., WilmesRiesenberg, M. R., & Wanner, B. L. (1996). Involvement of the sensor kinase EnvZ in the in vivo activation of the response-regulator PhoB by acetyl phosphate [Article]. Molecular Microbiology, 22(1), 135–147. <Go to ISI>://A1996VM85600014

23. Laub, M. T., & Goulian, M. (2007). Specificity in two-component signal transduction pathways. Annu Rev Genet, 41, 121–145. 10.1146/annurev.genet.41.042007.170548

24. Liu, W., & Hulett, F. M. (1997). *Bacillus subtilis* PhoP binds to the *phoB* tandem promoter exclusively within the phosphate starvation-inducible promoter [Article]. Journal of Bacteriology, 179(20), 6302–6310. <Go to ISI>://A1997YA93700012

25. Liu, W., Qi, Y., & Hulett, F. M. (1998). Sites internal to the coding regions of *phoA* and *pstS* bind PhoP and are required for full promoter activity [Article]. Molecular Microbiology, 28(1), 119–130. <Go to ISI>://000073122300011

26. Ludwig, H., Meinken, C., Matin, A., & Stulke, J. (2002). Insufficient expression of the *ilv-leu* operon encoding enzymes of branched-chain amino acid biosynthesis limits growth of a *Bacillus subtilis ccpA* mutant [Article]. Journal of Bacteriology, 184(18), 5174–5178. 10.1128/jb.184.18.5174-5178.2002

27. Lukat, G. S., McCleary, W. R., Stock, A. M., & Stock, J. B. (1992). Phosphorylation of Bacterial Response Regulator Proteins by low-molecular-weight Phospho-donors [Article]. Proceedings of the National Academy of Sciences of the United States of America, 89(2), 718–722. <Go to ISI>://A1992GZ69600055

28. McCleary, W. R., & Stock, J. B. (1994). Acetyl phosphate and the activation of 2-component response regulators. [Article]. Journal of Biological Chemistry, 269(50), 31567–31572. <Go to ISI>://A1994PX30300038

29. Nakano, M. M., & Hulett, F. M. (1997). Adaptation of *Bacillus subtilis* to oxygen limitation. FEMS Microbiol Lett, 157(1), 1–7. 10.1111/j.1574-6968.1997.tb12744.x

30. Nakano, M. M., Zhu, Y., Lacelle, M., Zhang, X., & Hulett, F. M. (2000). Interaction of ResD with regulatory regions of anaerobically induced genes in *Bacillus subtilis*. Molecular Microbiology, 37(5), 1198–1207. 10.1046/j.1365-2958.2000.02075.x

31. Ogura, M., Yamaguchi, H., Yoshida, K., Fujita, Y., & Tanaka, T. (2001). DNA microarray analysis of *Bacillus subtilis* DegU, ComA and PhoP regulons: an approach to comprehensive analysis of B.subtilis two- component regulatory systems. Nucleic Acids Research, 29(18), 3804–3813. 10.1093/nar/29.18.3804

32. Paul, S., Birkey, S., Liu, W., & Hulett, F. M. (2004). Autoinduction of Bacillus subtilis phoPR operon transcription results from enhanced transcription from E sigma(A)- and E sigma(E)-responsive promoters by phosphorylated PhoP [Article]. Journal of Bacteriology, 186(13), 4262–4275. <Go to ISI>://000222189500024

33. Puri-Taneja, A., Paul, S., Chen, Y. H., & Hulett, F. M. (2006). CcpA causes repression of the phoPR promoter through a novel transcription start site, P_A6_ [Article]. Journal of Bacteriology, 188(4), 1266–1278. 10.1128/jb.188.4.1266-1278.2006

34. Qi, Y., & (1998). Identification and Regulation of new PHO regulon genes in Bacillus subtilis [Doctor of Philosophy in Biological Sciences., University of Illinois at Chicago]. Chicago.

35. Schau, M., Eldakak, A., & Hulett, F. M. (2004). Terminal oxidases are essential to bypass the requirement for ResD for full Pho induction in *Bacillus subtilis* [Article]. Journal of Bacteriology, 186(24), 8424–8432. 10.1128/jb.186.24.8424-8432.2004

36. Shi, L., & Hulett, F. M. (1999). The cytoplasmic kinase domain of PhoR is sufficient for the low phosphate- inducible expression of Pho regulon genes in *Bacillus subtilis*. Molecular Microbiology, 31(1), 211–222. <Go to ISI>://WOS:000077954400020

37. Sonenshein, A. L. (2007). Control of key metabolic intersections in *Bacillus subtilis* [Review]. Nature Reviews Microbiology, 5(12), 917–927. 10.1038/nrmicro1772

38. Sun, G., Sharkova, E., Chesnut, R., Birkey, S., Duggan, M. F., Sorokin, A., … Hulett, F. M. (1996). Regulators of aerobic and anaerobic respiration in *Bacillus subtilis*. Journal of Bacteriology, 178(5), 1374–1385. 10.1128/jb.178.5.1374-1385.1996

39. Sun, G. F., Birkey, S. M., & Hulett, F. M. (1996). Three two-component signal-transduction systems interact for Pho regulation in *Bacillus subtilis* [Article]. Molecular Microbiology, 19(5), 941–948. <Go to ISI>://A1996TZ97400004

40. Trieucuot, P., Poyartsalmeron, C., Carlier, C., & Courvalin, P. (1990). Nucleotide sequence of the erythromycin resistance gene of the conjugative transposon TN1545 [Note]. Nucleic Acids Research, 18(12), 3660–3660. <Go to ISI>://A1990DL81500051

41. Wanner, B. L. (1992). Is cross regulation by phosphorylation of two-component response regulator proteins important in bacteria? Journal of Bacteriology, 174(7), 2053–2058. 10.1128/jb.174.7.2053-2058.1992

42. Williams, S. G., Cranenburgh, R. M., Weiss, A. M. E., Wrighton, C. J., Sherratt, D. J., & Hanak, J. A. J. (1998). Repressor titration: a novel system for selection and stable maintenance of recombinant plasmids [Article]. Nucleic Acids Research, 26(9), 2120–2124. <Go to ISI>://000073435300014

43. Yanischperron, C., Vieira, J., & Messing, J. (1985). Improved M13 phage cloning vecotrs and host strains- Nucletode sequence of the M13mp18 and pUC19 vectors [Article]. Gene, 33(1), 103–119. <Go to ISI>://A1985AEC6400002

44. Zhang, X., & Hulett, F. M. (2000). ResD signal transduction regulator of aerobic respiration in *Bacillus subtilis*: ctaA promoter regulation. Molecular Microbiology, 37(5), 1208–1219. 10.1046/j.1365-2958.2000.02076.x

